# Automated Generation of Cerebral Blood Flow Maps Using Deep Learning and Multiple Delay Arterial Spin-Labelled MRI

**DOI:** 10.1101/2021.06.04.446768

**Authors:** Nicholas J. Luciw, Zahra Shirzadi, Sandra E. Black, Maged Goubran, Bradley J. MacIntosh

## Abstract

The purpose of this work was to develop and evaluate a deep learning approach for estimation of cerebral blood flow (CBF) and arterial transit time (ATT) from multiple post-label delay (PLD) arterial spin-labelled (ASL) MRI. Six-PLD ASL MRI was acquired on a 1.5T or 3T system among 99 older males and females with and without cognitive impairment. We trained and compared two network architectures: standard feed-forward convolutional neural network (CNN) and U-Net. Mean absolute error (MAE) was evaluated between model estimates and ground truth obtained through conventional processing. The best-performing model was re-trained on inputs with missing PLDs to investigate generalizability to different PLD schedules. Relative to the CNN, the U-Net yielded lower MAE on training data. On test data, the U-Net MAE was 8.4±1.4 ml/100g/min for CBF and 0.22±0.09 s for ATT. Model uncertainty, estimated with Monte Carlo dropout, was associated with model error. Network estimates remained stable when tested on inputs with up to three missing PLD images. Mean processing times were: U-Net pipeline = 10.77s; ground truth pipeline = 10min 41s. These results suggest hemodynamic parameter estimation from 1.5T and 3T multi-PLD ASL MRI is feasible and fast with a deep learning image-generation approach.

## 1.0 Introduction

Arterial spin-labelled (ASL) MRI is becoming increasingly popular and available for examinations of cerebral blood flow (CBF) in dementia, stroke, and other neurologic and psychiatric diseases.^1^ While most CBF imaging modalities require an injected contrast agent, ASL is non-invasive and therefore uniquely suited for follow-up scans and vulnerable patient populations.

The rising availability of ASL in the clinic can be partially attributed to improvements in pulse sequence design of both single post-label delay (PLD) and, more recently, multi-PLD ASL.^2^ Multi-PLD ASL is desirable for clinical application because it enables imaging along different stages of the vascular network, thereby capturing dynamic information about inflowing blood.^3^ Such information offers the opportunity to quantify the arterial transit time (ATT) and mitigate the transit delay confounds evident in single-PLD ASL, as observed in clinical studies.^4–6^ Multiple PLDs, however, require cumbersome and time-consuming post-processing to synthesize CBF and ATT images, which limits the clinical adoption of multi-PLD ASL.^7^

Deep learning offers advantages for many stages of conventional image processing pipelines,^8^ as is the case for de-noising and artifact removal in single-PLD ASL processing.^9–12^ To date, however, the application of deep learning in ASL has been limited to single PLDs. Consideration of multiple PLDs, model uncertainty, and additional clinical populations are necessary to advance the application of deep learning methods in ASL.

In this study, we aimed to demonstrate that a deep learning-based processing pipeline is feasible for fast and accurate estimation of CBF and ATT from multi-PLD ASL. We performed three experiments using scans of older participants with and without cognitive impairment, for images acquired at either 1.5T or 3T. First, we trained two convolutional neural networks using each PLD as a separate input channel and performed cross-validation to select the best-performing architecture. Next, we examined the test-set performance of the best-performing network and estimated the association between network uncertainty and error. Finally, we investigated whether an alternative training scheme could improve model performance when tested on inputs with fewer PLD images, emulating alternative PLD schedules.

## 2.0 Methods

### 2.1 Participants

This is a retrospective study of 99 participants scanned between December 2009 and April 2014 at the Sunnybrook Research Institute (Toronto, ON). Research ethics approval was granted by the ethics board at Sunnybrook Research Institute. Participants were recruited through a neurology clinic at Sunnybrook Health Sciences Centre (detailed inclusion/exclusion criteria can be found at https://clinicaltrials.gov/ct2/show/NCT01800214). Participants were either cognitively unimpaired, had a mild cognitive impairment diagnosis, or had a dementia diagnosis.

### 2.2 Magnetic Resonance Imaging

Imaging was performed on a 1.5 Tesla GE Signa scanner (General Electric Medical Systems, Milwaukee, WI) for 50 participants and on a 3.0 Tesla GE Discovery 750 scanner for 49 participants. Both scanners were equipped with an 8-channel head array coil for signal reception. As detailed in the main text, the MRI protocol consisted of ASL and anatomical imaging, namely T1-weighted images. ASL MRI was nearly identical on the two scanners and included a 2D echo planar imaging acquisition, 64 × 64 × 17 matrix, 3.5 × 3.5 × 5.6 mm^3^ voxel resolution, TR/TE=3500-5000/17 ms, pseudo-continuous labelling (1500 ms label duration, six PLDs of 100, 500, 900, 1300, 1700, 2100 ms), and 12 control-label image pairs for each PLD. Labelling was performed using 1000 504μs Hanning-shaped 35° pulses. Additional ASL sequence details have been previously reported.^13^ A long-TR reference image for CBF quantification was also acquired at the beginning of each different PLD. Anatomical T1-weighted imaging was performed for registration and tissue segmentation. The anatomical 1.5 Tesla T1-weighted acquisition was an axial 3D SPGR with TR/TE= 35/5 ms, flip angle 35°, 256 × 256 matrix, 0.9 × 0.9 mm^2^ resolution, and 1.2-1.4 mm slice thickness, depending on participant head size. The 3.0 Tesla T1-weighted acquisition was an axial 3D FSPGR with TR/TE/TI: 8.1/3.2/650 msec, flip angle 8°, 256 × 256 matrix, 0.86 × 0.86 mm^2^ resolution, and 1.0 mm slice thickness.

### 2.3 Image Processing

#### 2.3.1 Ground truth pipeline

Ground truth parameter maps were estimated using a standard ASL MRI processing pipeline built with tools from the FMRIB Software Library (FSL). Motion correction was applied by registering the time series to the middle volume.^14^ CBF-weighted difference images were calculated via sinc subtraction of control and label volumes. Motion-corrupted difference images were discarded using the ENABLE quality control algorithm.^15,16^ Images were skull-stripped by applying a brain mask estimated from segmentation of the T1w image. CBF and ATT were estimated using the BASIL software, which incorporates ASL data from all acquired PLDs into a kinetic-model-based inference scheme.^17^ Resultant ATT images were in units of seconds and CBF images were converted to units of ml/100g/min using a standard calibration equation.^7^

#### 2.3.2 Proposed pipeline

Figure 1A shows the proposed pipeline schematic. First, difference images were calculated with sinc subtraction of raw control-label ASL volumes. The resultant images were averaged at each PLD, resulting in a six-channel input to the neural network. Input and ground truth image intensities were standardized by dividing each voxel by the 95^th^ percentile of intensities in the training sample. The network outputs were designed to be standardized CBF and ATT images that were subsequently scaled using the inverse of the standardization transform.

**Figure 1.**
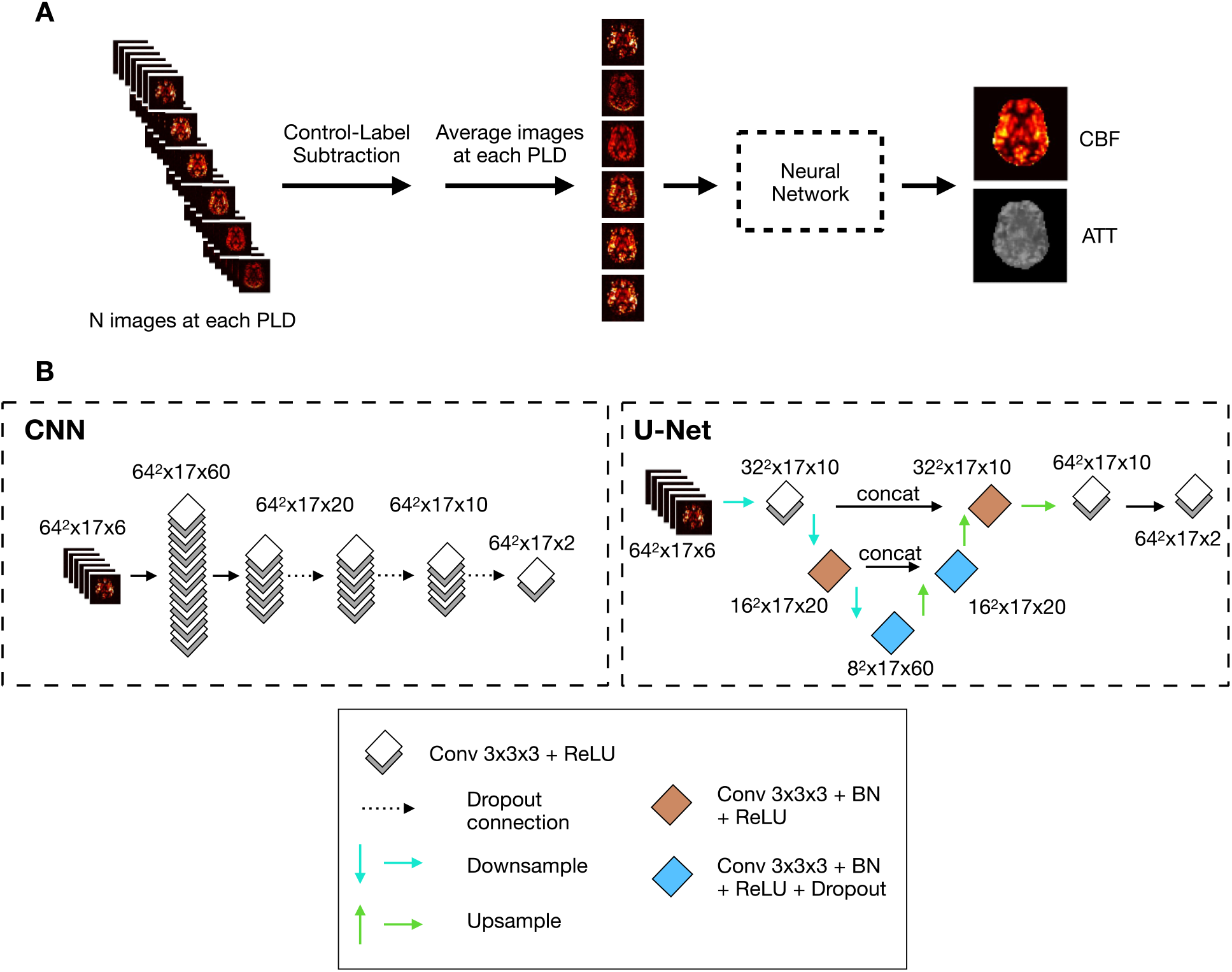
**A** Image processing pipeline. PLD=post-label delay. **B** Network architectures. Architectures are shown in dashed-line boxes, while the legend is shown in the solid-line box. ReLU=rectified linear unit; BN=batch normalization.

### 2.4 Network implementations

We implemented two distinct convolutional neural network architectures in Python 3.5 using the Keras API of the TensorFlow 2.2 library. As shown in Figure 1B, the two architectures were: 1) a 3D convolutional neural network (hereafter “CNN”) and 2) a U-Net.^18^ In both cases, trainable parameters were initialized with normally distributed random values.

For comparison of the two network architectures, the same six-image inputs were used (i.e. one 3D image per PLD), along with two output images (i.e. one 3D map for each of CBF and ATT). By design, each model varied in the number of trainable parameters.

The CNN consisted of five 3D convolutional layers with a 3 × 3 × 3 kernel followed by a ReLU activation function. The U-Net included seven convolutional layers with a 3 × 3 × 3 kernel, three of which down-sampled the image resolution (i.e. U-Net depth of 3) and three up-sampled the images back to the original resolution. Down and up-sampling was performed by a factor of two in the *x*-*y* plane using convolutions with stride 2.

### 2.5 Training details

We augmented each image (all input and output images) in the training set by applying three separate transformations: 1) mirroring in the *x-y* plane, 2) rotating by +20° in the *x-y* plane, and 3) rotating by −20° in the *x-y* plane. Therefore, the total dataset used for cross-validation was 316 image sets (comprising the original 79 times three augmentations for each).

Training of both networks was performed on a Linux system with an NVIDIA P100 Pascal GPU (NVIDIA, Santa Clara, California). For the CNN and U-Net, the loss function was the mean absolute error (MAE) adjusted to assign equal weighting to brain and background voxels, defined using the average of all images in the training set. Explicitly, the loss function was

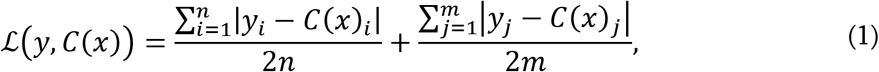

where *y* is the ground truth image, *C* is the neural network (CNN or U-Net), *x* is the input image, *n* is the number of brain voxels, and *m* is the number of background voxels. We used the Adam optimizer with an initial learning rate of 0.0005 which was reduced to 0.0001 after 100 epochs. Visual inspection of the loss curves was used to ensure plateau of the training and validation losses and to avoid overfitting. Each model was trained for 200 epochs during cross validation using a batch size of five image sets. The best performing model was trained for 600 epochs on the entire training set prior to testing, also using a batch size of five. The number of trainable parameters in each of the networks was: CNN=58,592; U-Net=88,792.

The best performing model architecture was selected based on the lowest aggregate (normalized CBF and ATT) MAE of grey matter voxels across all validation data in the cross validation.

### 2.6 Experiments

Prior to cross-validation, a random 80%/20% train/test split was employed. Splits were stratified by scanner magnetic field strength.

#### 2.6.1 Cross-validation for architecture selection

A five-fold cross-validation of the training set, stratified by scanner magnetic field strength, was used to select the best-performing network architecture. Performance was evaluated based on aggregate (CBF and ATT) mean absolute error (MAE) of grey matter voxels.

#### 2.6.2 Evaluation of model performance and model uncertainty estimation

We re-trained the architecture corresponding to the best-performing model on the entire training set. Performance was evaluated on the test set using MAE and the structural similarity index for both CBF and ATT. MAE was evaluated across all grey matter, all white matter, and five grey matter regions: frontal lobe, parietal lobe, occipital lobe, temporal lobe, and subcortical grey matter. Regions were identified using the MNI152 atlas and registered to each individual’s ASL image using FSL FLIRT, and using registration to the individual’s T1w image as an intermediate step.^19^

We used dropout (random deletion of network connections) during testing to produce non-deterministic network estimates.^20^ The dropout rate was set to 0.30. Two uncertainty heuristics were subsequently considered: standard deviation and coefficient of variation across 500 samples.

#### 2.6.3 Generalizing to cases with missing input images via curriculum training

We re-trained the model using a learning scheme in which one or more of the six input images were deliberately missing (i.e., image intensities set to zero). This strategy emulates changing the PLD schedule and reducing the scan duration. Examples with missing data were introduced during training in order of increasing difficulty, meaning more inputs missing and therefore less information provided (a strategy known as curriculum training).^21^ We removed inputs in a systematic outside-in fashion: first the 100ms PLD image was removed, then 2100ms, then 500ms.

Curriculum training was performed for 600 total epochs wherein the inputs with missing PLDs were introduced starting at epoch 200, after which the difficulty of training examples was increased (i.e., another PLD was removed) at epochs 300 and 400.

### 2.7 Processing Time Determination

Total processing time of both the proposed pipeline and the ground truth reference pipeline were estimated by timing the processing of a single dataset ten times.

### 2.8 Statistical Analyses

All data were tested for normality with the Shapiro-Wilk test. In the case of a significant result (*a priori* significance threshold = 0.05), non-parametric statistical tests were employed.

We performed a two-tailed *t*-test (*a priori* significance threshold = 0.05) to test for differences between field strengths in grey matter MAE of CBF and ATT (Wilcoxon signed-rank test was used for ATT due to evidence of non-normality).

The relationship between regional MAE in grey matter and the standard deviation uncertainty heuristic was investigated with a linear mixed-effects model with grey matter region as a random intercept term.

To evaluate whether curriculum training improves performance when three PLDs were removed from the input, a two-tailed Wilcoxon signed-rank test compared MAE of the curriculum-trained model with the model trained on only six-input images. Using a repeated-measures ANOVA, we tested for differences in CBF MAE of the curriculum-trained model between examples with zero, one, two, and three input images removed; in the case of ATT, we used the non-parametric equivalent of repeated-measures ANOVA, the Friedman test, because there was evidence for non-normality of ATT MAE.

### 2.9 Code availability

Python code used to train the models, evaluate performance, and generate the figures presented in this manuscript will be available at *https://github.com/nluciw/Deep-learning_for_mPLD-ASL*. The TensorFlow-based models and weights will also be available.

## 3.0 Results

### 3.1 Participant demographics

Demographics and basic clinical information of the 99 participants are shown in Table 1. Scanning was performed at 1.5 T for 50 participants and at 3.0 T for 49 participants. 23 participants were cognitively unimpaired, 17 had a mild cognitive impairment diagnosis, and 59 had a dementia diagnosis.

**Table 1.** Participant demographics. Additional detailed diagnostic information is provided for the group with a dementia diagnosis. SD=standard deviation.

|  | 1.5T Images<br>(N = 50) | 3.0T Images<br>(N = 49) |
| --- | --- | --- |
| **Demographics** |  |  |
| Age, mean (SD) | 73.1 (8.8) | 73.6 (8.1) |
| Females, n (%) | 27 (54.0) | 30 (61.2) |
| **Dementia Diagnosis** |  |  |
| Alzheimer's Disease | 15 | 23 |
| Cortical Basal Syndrome | 3 | 0 |
| Dementia with Lewy Bodies | 3 | 0 |
| Frontotemporal Dementia | 7 | 4 |
| Vascular Dementia | 1 | 0 |
| Cortical Basal Syndrome / Progressive Supranuclear Palsy | 1 | 0 |
| Dementia with Lewy Bodies / Parkinson's Disease Dementia | 1 | 0 |
| Alzheimer's Disease / Frontotemporal Dementia | 1 | 0 |
| Mild Cognitive Impairment | 16 | 1 |
| Cognitively unimpaired | 2 | 21 |

### 3.1 Selection of network architecture: cross-validation on training data

The MAE of CBF estimates across five-fold cross-validation were: CNN = 10.3 ± 5.0 ml/100g/min and U-Net = 9.2 ± 1.7 ml/100g/min. The MAE of ATT estimates were: CNN = 0.30 ± 0.18 s and U-Net = 0.27 ± 0.18 s. Because the U-Net produced estimates with the lowest MAE, we chose the U-Net as the best-performing architecture to consider for the next experiments.

### 3.2 Test set performance of the U-Net

The performance of the U-Net on the test set is shown in Figure 2 and Table 2. Average (mean ± standard deviation) grey matter MAE for CBF and ATT estimates was 8.4 ± 1.4 ml/100g/min and 0.22 ± 0.09 s. Average grey matter MAE was lower for CBF (two-tailed *t*-test: *t*=−3.04, *p*=0.014), but not ATT (two-tailed Wilcoxon signed-rank test: *W*=25.0, *p*=0.799), in the scans acquired at 1.5T compared to 3T. Representative high and low MAE network estimates and corresponding ground truth images are shown in Figure 3.

**Figure 2.**
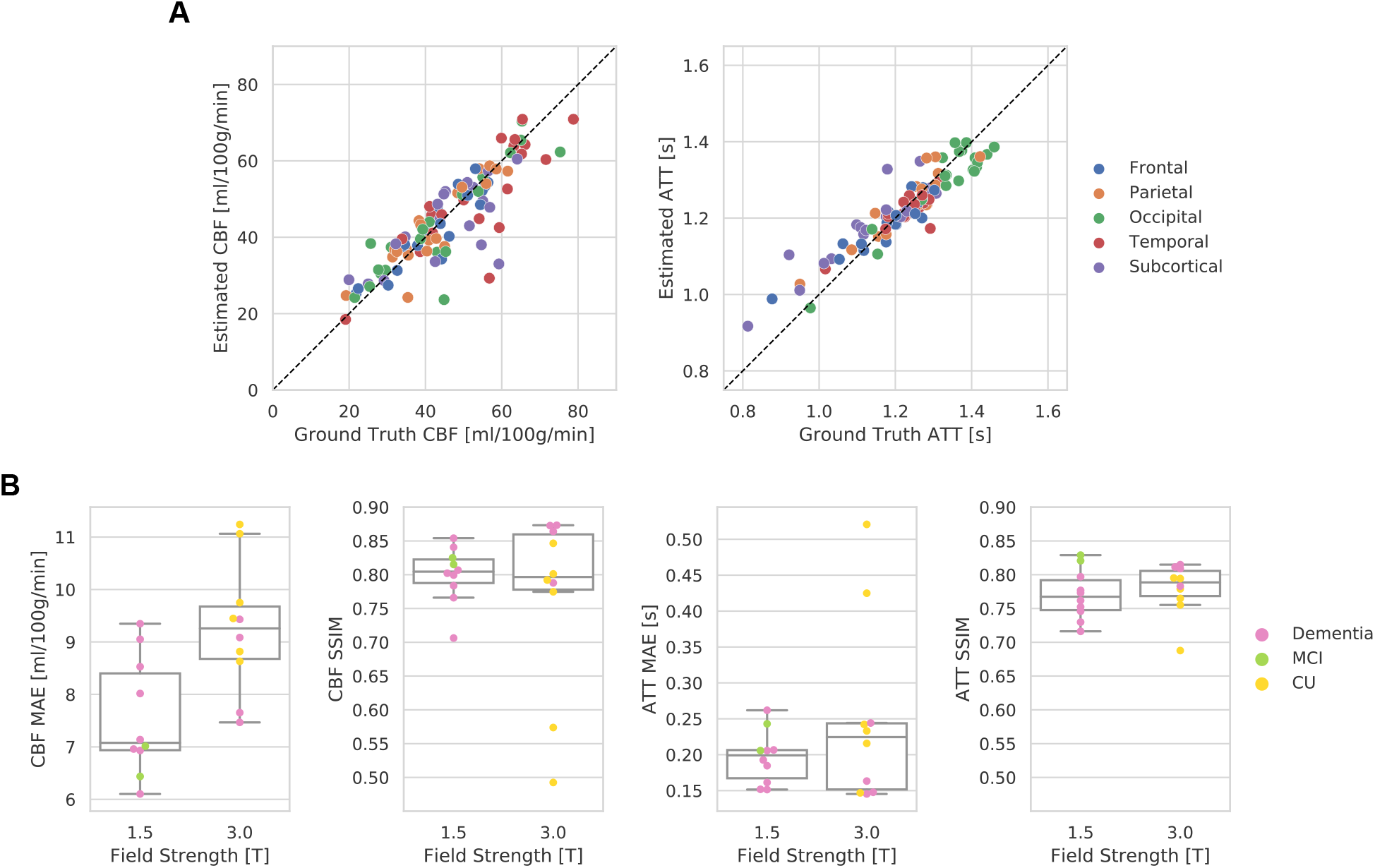
Test set performance. **A** Association between network estimate and reference images averaged across five grey matter regions (resulting in five data points per participant). Colours denote grey matter region. **B** Mean absolute error (MAE) and structural similarity (SSIM) index between network estimate and ground truth. Samples are stratified by scanner field strength. Colours denote clinical status. MCI=mild cognitive impairment; CU=cognitively unimpaired.

**Table 2.** Test set (n=20) performance characteristics. For reference, average ground truth grey matter CBF and ATT across the test set was 51.1 ± 12.1 ml/100g/min and 1.21 ± 0.08 s. std=standard deviation.

|  | **CBF (mean ± std)**<br>[ml / 100g / min] | **ATT (mean ± std)**<br>[s] |
| --- | --- | --- |
| **Mean absolute error** |  |  |
| Grey matter | $8.4 \pm 1.4$ | $0.22 \pm 0.09$ |
| Frontal lobe | $8.3 \pm 2.1$ | $0.18 \pm 0.07$ |
| Occipital lobe | $8.2 \pm 1.5$ | $0.25 \pm 0.11$ |
| Parietal lobe | $7.6 \pm 1.4$ | $0.16 \pm 0.07$ |
| Temporal lobe | $9.1 \pm 1.8$ | $0.30 \pm 0.15$ |
| Subcortical | $7.5 \pm 3.0$ | $0.16 \pm 0.13$ |
| White matter | $7.5 \pm 1.9$ | $0.16 \pm 0.07$ |
| **Structural similarity**<br>(entire image; unitless) | $0.78 \pm 0.10$ | $0.77 \pm 0.04$ |

**Figure 3.**
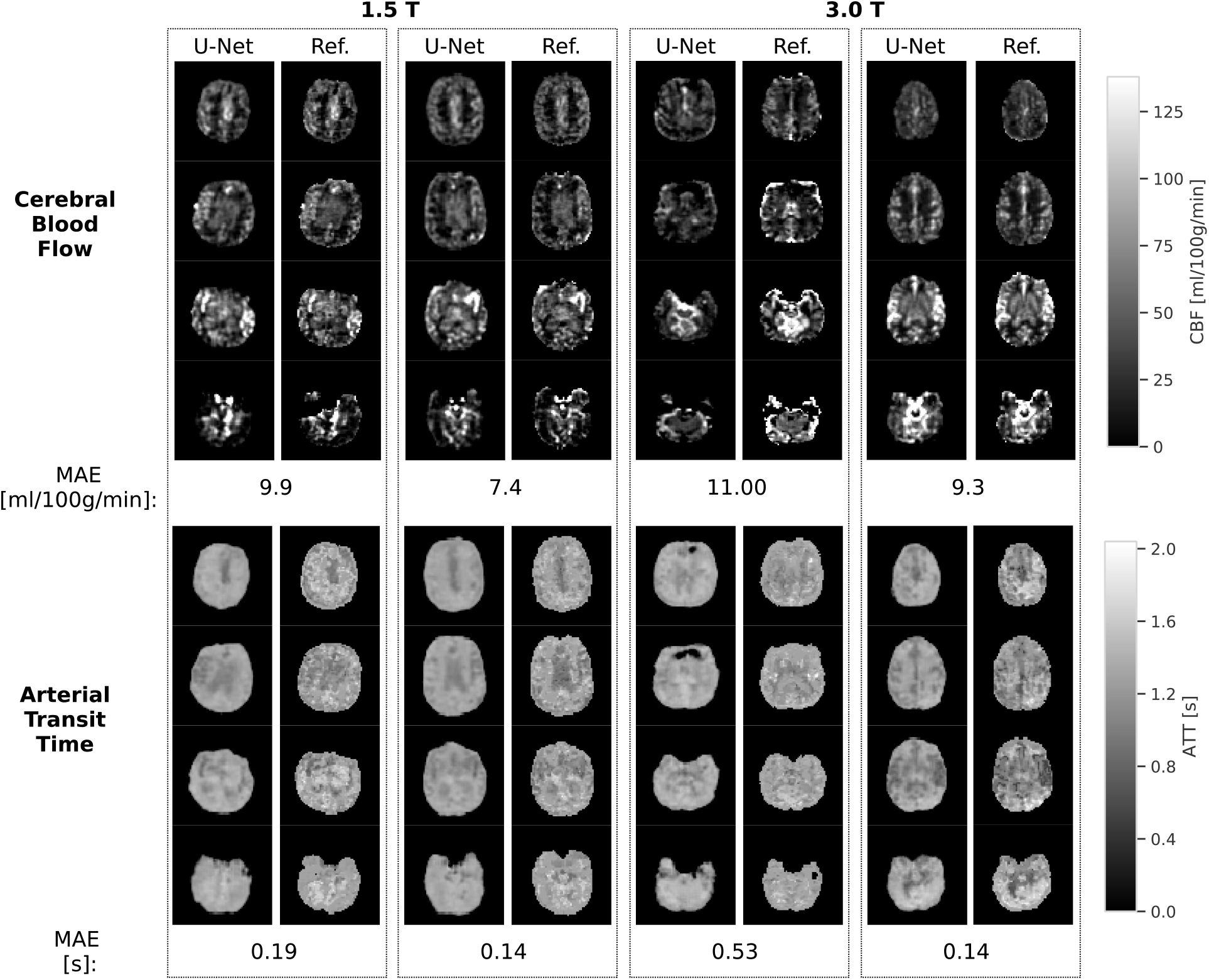
High-quality and low-quality examples of CBF and ATT estimates for both 1.5T and 3T scans. Each panel displays four axial slices from the same image (top: most superior, bottom: most inferior). Each pair of panels displays the network estimate and the reference ground truth on the left and right, respectively. Each column displays images from the same individual. Ref.=reference ground truth image, MAE=mean absolute error.

### 3.3 Network uncertainty estimation

Figures 4A and 4B show the network uncertainty heuristics and mean absolute error for CBF and ATT. Across individuals, a linear mixed effects model revealed an association between uncertainty and MAE of both CBF (*β*=0.16, *t*=2.64, *p*=0.0097) and ATT (*β*=0.048, *t*=3.43, *p*=0.0009). Representative uncertainty maps for high and low-quality CBF estimates are shown in Figure 4C and 4D, respectively.

**Figure 4.**
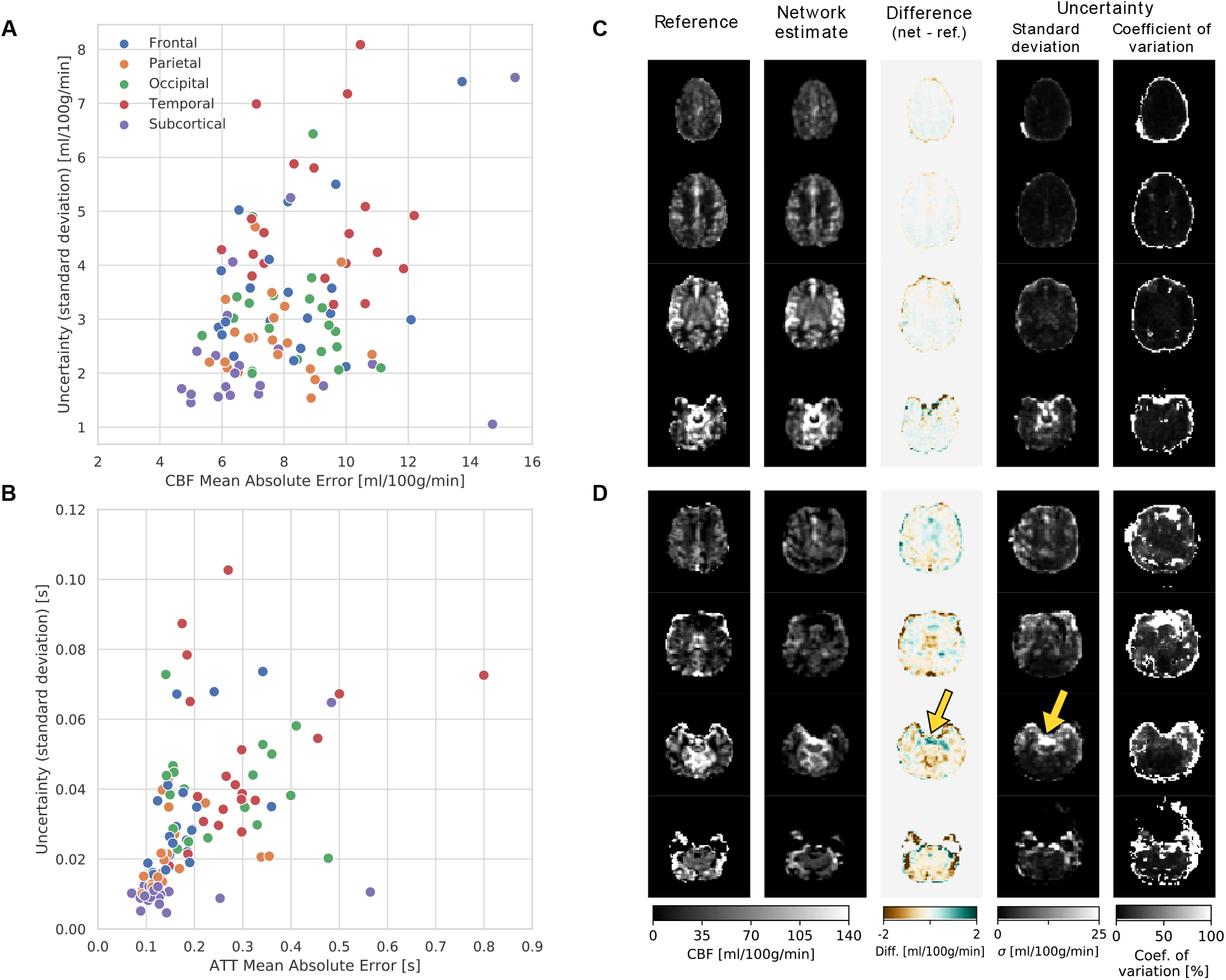
Network uncertainty heuristics estimated through Monte Carlo dropout. **A** Association of standard deviation with mean absolute error of regional grey matter CBF estimates in each test set participant. Colours denote grey matter region. **B** Association of network standard deviation with mean absolute error of ATT estimates. **C** High-quality 3T CBF estimate alongside reference ground truth, difference between estimate and ground truth reference, and uncertainty estimates. **D** Low-quality 3T CBF estimate alongside reference ground truth, difference between estimate and ground truth reference, and uncertainty estimates. Yellow arrows highlight an example of an area where high uncertainty overlaps with high error.

### 3.4 Curriculum training for parameter estimation with less than six input images

MAE as a function of missing ASL data is shown in Figure 5. Two models are depicted: one trained on examples with missing input data (i.e. curriculum-trained) and one trained with only the full input data. The curriculum-trained model had significantly lower CBF MAE relative to the model trained on full inputs when considering inputs with three images removed (two-tailed Wilcoxon signed-rank test: *W*<0.1, *p*=8.86×10^−5^). Repeated-measures ANOVA revealed no differences in CBF MAE of the curriculum-trained model across the schedules with zero, one, two, and three input images removed (*F*=1.70, *p*=0.177).

**Figure 5.**
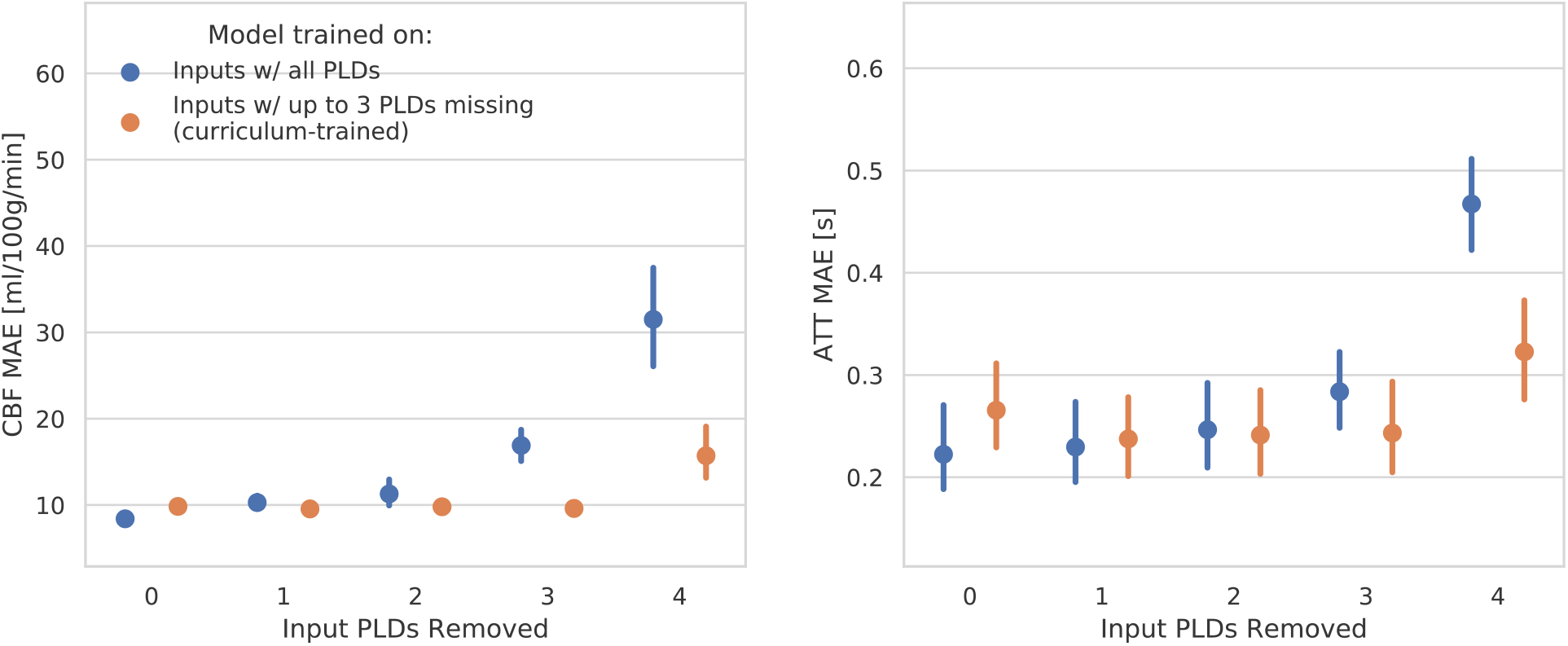
Model performance when PLDs are removed from the input, emulating a different PLD schedule. Performance of two models is shown: 1) the model trained on examples with all six images as input; 2) the curriculum-trained model, trained on examples with up to three input images missing. Translucent points show individuals’ data, large solid circles show mean, and bars show 95% confidence intervals. MAE=mean absolute error; PLD=post-label delay.

For ATT estimates, the curriculum-trained model similarly had significantly lower MAE relative to the model trained using full inputs when considering inputs with three images removed (two-tailed t-test *t*=−5.67, *p*=1.8×10^−5^). There were significant differences in ATT MAE of the curriculum-trained model across the schedules with zero, one, two, and three input images removed (Friedman test: *F*=19.92, *p*=0.0002).

### 3.5 Processing times for U-Net and ground truth pipelines

The total processing time of the U-Net approach was 10.77 ± 0.19 s. The U-Net model alone took 2.44 ± 0.03 s. In comparison, the total processing time of the ground truth calculation was 10 min 41 s ± 40.8 s (of which approximately 4 mins was for tissue segmentation used by the ENABLE quality control algorithm).

## 4.0 Discussion

In this study, we trained deep convolutional neural networks to estimate cerebral blood flow and arterial transit time maps from multi-PLD ASL MRI. The U-Net modestly outperformed the standard CNN with average CBF estimation error of 8.4 ± 1.4 ml/100g/min on the test set. U-Net uncertainty estimates were associated with MAE of the network-generated CBF and ATT maps. Lastly, through a curriculum learning scheme, we trained the U-Net to improve its performance when fewer image inputs were provided. The U-Net-based pipeline was substantially faster than the ground truth pipeline, a crucial consideration in time-constrained clinical settings.

Overall, the average test set CBF error of the U-Net model was comparable to the variability that tends to occur because of different processing choices in a standard ASL pipeline, reported to be between 5-10 ml/100 g/min.^22^ This finding suggests our proposed pipeline is feasible for CBF estimation and there is no data quality penalty despite substantive savings in processing time.

The U-Net outperformance of the CNN during cross-validation was modest and may be sample-specific or related to the number of trainable parameters and architecture design. Further hyperparameter optimization may produce different results.

In the test set, we observed a significant effect of scanner field strength on network CBF MAE; somewhat unexpectedly, 1.5T data had lower MAE than 3T data, which may reflect two considerations. First, with a lower signal-to-noise ratio, 1.5T data could permit the network to learn average noise features. Second, the participant demographics were different for the two scanners, with more cognitively unimpaired individuals scanned at 3T. We also observed two outliers in this 3T cohort, based on low structural similarity index. These outliers were less prominent based on the MAE metric, which may reflect both the choice of MAE as the training loss function and the fundamental differences between MAE, which assesses differences between each voxel, and structural similarity, which is a whole-image summary metric.

Estimating model uncertainty provided insight on regional model performance without the need for ground truth comparison. Expedient access to uncertainty information is pertinent in clinical deployment of deep learning methods: uncertainty can offer quantification of image quality and is a potential avenue for automated quality control. Indeed, the standard deviation-based uncertainty heuristic was associated with both CBF and ATT errors in grey matter. As a secondary metric, we also evaluated uncertainty based on the coefficient of variation, which displayed qualitatively distinct contrast relative to standard deviation. This was likely due to association between standard deviation and CBF; therefore, the coefficient of variation, which is the standard deviation relative to CBF, may be differently sensitive to errors.

By training the network on inputs with missing PLDs from the six-PLD acquisition, we demonstrated preliminary evidence that the network may generalize to accommodate different acquisitions, which is desirable to allow patient-specific PLD schedules and to reduce scan duration. As expected, the model performed better on three-PLD inputs after it trained on examples with a three-PLD input. Notably, this curriculum-trained model exhibited no performance decrements in CBF estimates across inputs with up to three missing PLDs. Therefore, a single model may be used in a setting where different PLD schedules are acquired, without the need for re-training. Conversely, considering ATT estimates, the curriculum-trained model performed better using inputs with three PLDs compared to six, five, or four PLDs; further attention to the learning scheme may be warranted to improve ATT estimation.

Study limitations are discussed herein. First, the models do not explicitly incorporate tissue equilibrium magnetization, which is vital for absolute quantification of CBF. In the current study, the model may learn to scale appropriately because there are only two scanners; generalizability would require additional investigation of a broader set of pulse sequences and scanners. Second, although we implemented two network architectures, we did not perform exhaustive hyperparameter tuning. Thus, the results of the current study provide a benchmark for future studies. Third, the cognitive neurology sample was relatively small, which limits the investigation of sources of performance variability and sensitivity to diagnosis. Finally, the imaging sequence studied here is one implementation of multi-PLD ASL and, in the absence of a standardized protocol, future work is needed to expand on pulse sequence parameters.

In this study, we demonstrated the feasibility of using a deep neural network to estimate CBF and ATT from multi-PLD ASL MRI. We highlighted an avenue for quality control via network uncertainty heuristics and demonstrated the potential to generalize to a range of multi-PLD schedules. By simplifying and improving efficiency of the ASL workflow, this deep learning-based pipeline is a step toward wider adoption of multi-PLD ASL MRI.

## 5.0 Acknowledgments

We thank the individuals and their families for participating in this study. Additional thanks to the staff of the L.C. Campbell Neurology Unit at Sunnybrook Research Institute in Toronto, Canada.

## 6.0 Author contributions

Conception and design: NJL, ZS, MG, BJM. Data collection: SEB, BJM. Analysis and interpretation: NJL, BJM. Manuscript writing: NJL, BJM. Manuscript revision: NJL, ZS, SEB, MG, BJM. All authors approve the final version of this manuscript.

